# Frontoparietal functional dedifferentiation during naturalistic movie watching among older adults at risk of emotional vulnerability

**DOI:** 10.1101/2025.05.12.653474

**Authors:** Shuer Ye, Arjun Dave, Alireza Salami, Maryam Ziaei

**Affiliations:** Kavli Institute for Systems Neuroscience, Norwegian University of Science and Technology, Trondheim, Norway; Umeå Center for Functional Brain Imaging (UFBI), Umeå University, Umeå, Sweden; Department of Medical and Translational Biology, Umeå University, Umeå, Sweden; Wallenberg Center for Molecular Medicine, Umeå University, Umeå, Sweden; Aging Research Center, Karolinska Institutet & Stockholm University, Stockholm, Sweden; K.G. Jebsen Centre for Alzheimer’s disease, Norwegian University of Science and Technology, Trondheim, Norway

**Keywords:** movie-fMRI, emotion regulation, depression, frontoparietal network, gradient dispersion

## Abstract

Functional dedifferentiation, a hallmark of brain aging particularly evident within the frontoparietal network (FPN), has been extensively investigated in the context of cognitive decline, yet its implications for late-life mental health remain poorly understood. Leveraging naturalistic fMRI combined with gradient mapping techniques, the present study investigated FPN functional dedifferentiation—quantified by functional dispersion of FPN in the multidimension gradient manifold—during real-life emotional experiences and its link to affective outcomes. Here, we estimated functional dispersion during naturalistic movie watching in both younger (N=72, 34 female, 19-36 yrs) and older (N=68, 36 female, 65-82 yrs) adult groups with 7T MRI scanner and assessed their emotion regulation difficulties, anxiety, and depression symptoms as indicators of mental health status. The results demonstrated that greater FPN dispersion (i.e., more dissimilar connectivity) was linked to increased depressive symptoms in older adults and highlighted emotion regulation difficulties as a full mediator of this relationship. Moreover, FPN dispersion could distinguish emotionally resilient from vulnerable older individuals. These findings suggest that functional dedifferentiation of the FPN during ecologically valid emotional context constitutes a promising neural signature of affective vulnerability in older adults. By bridging age-related functional dedifferentiation to real-world emotional scenario, this work underscores the translational value of naturalistic paradigms in geriatric psychiatry and identifies potential intervention targets aimed at enhancing FPN specificity to promote mental health in aging population.

## 1. Introduction

The rapid aging of the global population has made the health challenges of older adults a critical public health priority (Dall et al., 2017). Among these challenges, mental health issues have become a major concern for researchers and clinicians (Dogra et al., 2022; Petrova & Khvostikova, 2021; Reynolds 3rd et al., 2022). Anxiety and depression, which frequently co-occur, are among the most common mental disorders in older adults (Bobo et al., 2022). Epidemiological studies estimate that approximately 35% of older adults experience clinically significant symptoms of depression or anxiety (Cai et al., 2023; Y. Liu et al., 2023; Zenebe et al., 2021). These mental health disorders not only exacerbate age-related cognitive decline and physical frailty but also increase the risk of all-cause mortality, thus posing significant barriers to healthy aging (X. Gao et al., 2023; Meier et al., 2016; Zannas, 2024).

Emotional regulation plays a crucial role in mental well-being across the lifespan, influencing how individuals manage stress, process negative emotions, and adapt to adversity (Gross, 2015; Miu et al., 2022). Effective emotional regulation strategies, such as cognitive reappraisal and problem-solving, contribute to psychological resilience and overall mental health (Uusberg et al., 2019). Conversely, difficulties in emotion regulation may contribute to increased vulnerability to mental disorders in both younger (Lincoln et al., 2022) and older groups (J. Sun et al., 2020). Research suggests that individuals with depression and anxiety exhibit difficulties in regulating negative emotions, leading to persistent distress and impaired social functioning (Chen & Bonanno, 2021; Martin & Dahlen, 2005). Mounting evidence indicates that emotional regulation is a complex cognitive process that relies on the coordinated activity of large-scale brain networks, among which the frontoparietal network (FPN) plays a crucial role (Almdahl et al., 2023; X. Cui et al., 2024; Morawetz et al., 2020). The FPN, encompassing regions of the prefrontal and parietal cortices, is primarily involved in cognitive control, goal-directed behavior, and attentional modulation. This network enables individuals to regulate emotions by integrating cognitive processes to effectively modulate affective responses (Bo et al., 2024; W. Li et al., 2021). Importantly, disruptions in the functions of FPN have been observed in individuals with affective disorders, highlighting (Belleau et al., 2023; Ding et al., 2024; Y. Gao et al., 2023; Ren et al., 2025; Zhang et al., 2025). For example, reduced functional connectivity within the FPN has been observed in patients with major depressive disorder (MDD), which is linked to rumination symptoms and suggests impaired goal-directed attentional control (Kaiser et al., 2015). Moreover, topological analysis of brain networks in MDD patients has also revealed reduced network segregation between the FPN and the default mode network (DMN), indicating alteration in functional network boundaries that contribute to emotional dysregulation (Lan et al., 2022).

The aging brain undergoes a reorganization characterized by neural dedifferentiation (Koen et al., 2020; Koen & Rugg, 2019), representing a progressive shift from functional specialization to integration, which is particularly pronounced within FPN (Cox, 2024; Rajah & D’Esposito, 2005; Shen et al., 2025). Neuroimaging studies have revealed that age-related dedifferentiation manifests as reduced modular segregation and increased between-network interactions (Raykov et al., 2024; Wu et al., 2023), with the FPN showing a significant decrease in network specificity compared to younger counterparts (Deery et al., 2023; X. Liu et al., 2023). Previous evidence has indicated functional dedifferentiation, reflecting an imbalance between segregation and integration as one potential mechanism of cognitive decline in aging (Bowman et al., 2019; J. Park et al., 2010; Varangis et al., 2019). While extensive research has established the consequences of functional dedifferentiation for cognitive processes in aging (Dennis & Cabeza, 2011; Yassa et al., 2011), its implications for affective states and psychological well-being remain largely underexplored. Specifically, given the role of FPN in cognitive control, its dedifferentiation may not only impair higher-order cognition but also disrupt emotion regulation, increasing susceptibility to affective disorders (Bo et al., 2024; Morawetz, 2020). Emerging evidence suggests that decreased functional connectivity within the FPN, reflecting its organizational dedifferentiation, is linked with late-life depression (H. Li et al., 2020). Thus, FPN dedifferentiation may serve as a neurobiological marker of increased vulnerability to mental health disorders in older adults (Almdahl et al., 2023; H. He et al., 2023). Characterizing this neurobiological marker is crucial for preserving mental health in older adults, as it may aid in the early identification of individuals at risk for emotional disorders (Tadayonnejad & Ajilore, 2014). A deeper understanding of FPN dedifferentiation could inform targeted interventions aimed at enhancing cognitive control over emotions, ultimately mitigating the impact of age-related declines in emotional regulation and reducing the prevalence of late-life depression and anxiety (Wen et al., 2022).

While traditional neuroimaging investigations of network dedifferentiation in aging have predominantly relied on task-based and resting-state paradigms, these approaches inherently limit ecological validity when examining complex, real-world dynamics (Finn, 2021). The emergence of naturalistic paradigms addresses this critical limitation by immersing participants in ecologically rich stimuli (e.g., emotionally charged movies) that preserve contextual embedding while minimizing experimental constraints (Kringelbach et al., 2023). Therefore, leveraging naturalistic paradigms (e.g., movie-fMRI), resembling real-life scenarios, provides a much deeper understanding of how the brain processes and regulates emotions in everyday life (Jääskeläinen et al., 2021). Studies have demonstrated that movie-fMRI allows for the precise characterization of individual differences in cognitive functions, emotional responses, social cognition, and personality traits by capturing stimuli-dependent brain dynamics in a real-world context (Esmaeili et al., 2025; Finn & Bandettini, 2021; Lee Masson et al., 2024; Meer et al., 2020). Given that dedifferentiation is thought to reflect a loss of functional specialization, movie-fMRI provides an ideal platform for examining how network organization dynamically adapts to emotionally salient stimuli.

Moreover, traditional connectivity-based analyses may overlook subtle, large-scale organizational changes associated with network organization including the dedifferentiation (Bernhardt et al., 2022; Lioi et al., 2021). To address this, we employed gradient analysis—a novel data-driven multidimensional framework approach that maps high-dimensional functional connectivity patterns into a low-dimensional manifold space—to specifically quantify the dedifferentiation of the FPN from macroscale brain organization perspective (Margulies et al., 2016). By calculating the functional dispersion of FPN regions along the main gradients that explain a large proportion of the variance in functional connectivity profiles, we defined dedifferentiation as an increased spatial Euclidean distance among FPN regions in the manifolds, reflecting greater heterogeneity in connectivity profiles and diminished functional specialization within the network (J. Li et al., 2024). Studies have shown that gradient dispersion, as a complement to the conventional network connectivity approach, provides valuable insights in detecting age-related brain changes (Bethlehem et al., 2020; J. Li et al., 2024), and it has been instrumental in clinical investigations such as depression(Pasquini et al., 2023; L. Sun et al., 2025), Alzheimer’s disease (Y. He et al., 2023), and bipolar disorder (Lei et al., 2023).

The aim of the current study was to investigate whether dedifferentiation of the FPN is linked to psychological well-being in aging, and to elucidate its underlying neuropsychological mechanisms by combining movie-fMRI and gradient analysis. We analyzed data from the Trondheim Aging Brain Study (TABS) cohort, in which younger and older adults underwent ultra-high field 7T fMRI scanning while viewing emotionally neutral and negative movie clips. Participants also completed self-reported questionnaires assessing emotion regulation difficulty, anxiety symptoms, and depressive symptoms. Functional dedifferentiation of the FPN was quantified by calculating its dispersion in gradient space and its association with emotion regulation difficulties, depression, and anxiety within younger and older adult groups. Furthermore, we employed latent profile analysis (LPA), a data-driven approach that detects hidden subgroups based on patterns within continuous observations, to uncover subgroups with distinct emotional well-being profiles and assess differences in FPN dedifferentiation across these subgroups. We hypothesized that: (1) FPN functional dispersion would be positively associated with symptoms of anxiety/depression in older adults, while no significant association is anticipated in younger adults (C.f. Brosnan et al., 2022); (2) Emotion regulation difficulties is expected to mediate the relationship between FPN dedifferentiation and affective dysregulation symptoms in the older group(Sharma et al., 2023); (3) and finally, FPN dispersion will enable the identification of individuals who are vulnerable to affective disorders in aging populations.

## 2. Methods

### 2.1 Participants

The data analyzed for the current study were collected as part of the TABS, focusing on emotional processing in aging. This study consists of data from 80 younger adults and 80 older adults recruited from local communities via advertisement. All participants were healthy individuals with normal auditory function and normal or corrected-to-normal visual acuity, and none had contraindications for MRI and a history of psychiatric or neurological disorders. After excluding technical issues, incomplete data, and excessive movement (i.e., mean frame displacement [FD] > 0.3 mm), data from 72 younger adults (34 females, mean age: 25.70 ± 4.33 yrs, age range: 19-36 yrs) and 68 older adults (36 females, mean age: 70.90 ± 3.83 yrs, age range:65-82 yrs) were included in the formal analyses. All participants completed two sessions, one imaging and one behavioral session, provided consent forms and were compensated with a gift card valued at 500 NOK for their participation. The study was conducted in accordance with the Declaration of Helsinki and had received ethical approval from the regional committees for medical and health research ethics.

### 2.2 Movie clips

This study utilized two movie clips during fMRI scanning: a neutral and a negative clip. The neutral clip, titled Pottery, depicts two women engaged in pottery-making and has a duration of 480 seconds. The negative clip, titled Curve, portrays a woman struggling to avoid falling into an abyss on a slippery cliff and lasts 494 seconds. The two movie clips were selected following a pilot study in which participants’ emotion assessment, emotional arousal, and valence ratings were collected for four clips. These selected two showed high validity as neutral and negative clips and thus were included in the imaging experiment. More details of these two clips can be found in our previous work (Ye et al., 2024), in which we demonstrated that Curve contains emotionally salient content and effectively inducing stress among participants. In contrast, Pottery serves as an effective control, as it elicits minimal emotional responses based on self-report responses. Therefore, in the present study, we primarily focused on the negative movie clip as our primary condition, using the neutral movie clip as a control condition.

### 2.3 Neuropsychological assessments

As part of the TABS, a series of cognitive and behavioral measurements including self-report questionnaires were administered outside the MRI scanner to assess executive functioning, mental well-being, emotion regulation capacity, and empathic tendencies (for more information regarding these measures see Dave et al., for details, Dave et al., 2024). While the full battery encompassed multiple psychological domains, the present investigation specifically prioritizes affective disorder (i.e., depression and anxiety) symptoms and emotion regulation as core variables of interest, given their strong link with emotional resilience and mental well-being shown in previous studies (Maddock, 2024; Zhi & Derakhshan, 2024). Consequently, our analyses focus on the Difficulties in Emotion Regulation Scale (DERS) and the Hospital Anxiety and Depression Scale (HADS).

The DERS was administered to assess multidimensional deficits in emotion regulation capacity(Gratz & Roemer, 2004). This 36-item self-report questionnaire uses a 5-point Likert scale (1=“almost never” to 5=“almost always”), with total scores ranging from 36 to 180, where higher scores indicate greater emotion dysregulation. The DERS comprises six subscales: i) nonacceptance of emotional responses, ii) difficulties engaging in goal-directed behavior, iii) impulse control difficulties, iv) lack of emotional awareness, v) limited access to emotion regulation strategies, and vi) lack of emotional clarity. In the present study, we use the sum of the items (i.e., total DERS score) to indicate the level of emotion regulation difficulties of participants, with a higher score indicating greater emotion regulation difficulty. Cronbach’s alpha of this scale was 0.884 in the current sample.

The HADS was administered to screen for anxiety and depression symptoms in non-psychiatric populations (Zigmond & Snaith, 1983). This 14-item self-report questionnaire consists of two subscales: Anxiety and Depression, each containing 7 items. Responses are recorded on a 4-point Likert scale (0=“not at all” to 3=“most of the time”), with subscale scores ranging from 0 to 21. Higher scores indicate greater symptom severity. Cronbach’s alpha was 0.792 and 0.789 for depression and anxiety subscale, respectively.

### 2.4 Procedure

Prior to the MRI session, participants were instructed on the procedure and experiment through both verbal and written instructions. During the MRI scan, participants first completed an 8-minute anatomical scan, followed by a movie-fMRI scan. They viewed a neutral movie clip, succeeded by a negative movie clip, and were instructed to rate their emotional valence and arousal levels before and after each clip using a 9-point Self-Assessment Manikin scale (for emotional valence rating, score 1=“very unpleasant” to 9=“very pleasant“; for emotional arousal, score 1=“very calm” to 9=“very excited”) (Bradley & Lang, 1994). Participants were instructed to watch the videos as naturally as possible while maintaining minimal head movement during the scanning session. To reduce head motion, foam padding was utilized. Auditory stimuli from the videos were delivered via MRI-compatible earphones (BOLDfonic, Cambridge Research Systems Ltd) and sound volumes were adjusted per individual prior to starting the session. Behavioral session was completed within three days from the MRI session.

### 2.5 Imaging acquisition and preprocessing

MRI scanning was conducted on a 7T MRI Siemens MAGNETOM Terra scanner equipped with a 32-channel head coil at Norwegian 7T MR Center, St Olav Hospital. Functional images were acquired using a multi-band accelerated echo-planar imaging sequence (92 interleaved slices, multi-band acceleration factorlJ=lJ8, voxel size = 1.25 × 1.25 ×1.25 mm, repetition time = 2000 ms, echo time = 19 ms, matrix size = 160 × 160 mm, field of view = 200 mm, slice thickness = 1.25 mm, and flip angle = 80°), resulting in 243 volumes for the neutral movie and 250 volumes for the negative movie. Anatomical images were acquired using a MP2RAGE sequence with 224 sagittal slices (voxel size = 0.8 × 0.8 × 0.8 mm, echo time = 1.99 ms, repetition time = 4300 ms, inversion time 1 = 840 ms, inversion time 2 = 2370 ms, flip angle = 5°/6° and slice thickness = 0.75 mm).

The functional imaging data were initially preprocessed using the standard pipeline in fMRIprep version 22.0.2(Esteban et al., 2019), followed by post-preprocessing with XCP-D version 0.3.053(Mehta et al., 2024). Preprocessing included removal of the first 3 volumes, motion and slice timing correction, co-registration, normalization to MNI space with 2 mm resolution, nuisance regression, band-pass filtering, and spatial smoothing with 4mm kernel size (details of analyses and codes are reported in **Supplementary Material**).

### 2.6 Gradient analysis

To estimate functional gradients, we first constructed individual functional connectomes for each participant. This was accomplished by parcellating the brain into 1,000 spatially independent regions using the Schaefer 1000-parcel atlas(**Fig.1C**) (Schaefer et al., 2018). The parcels can be assigned into 7 predefined functional networks: visual network, somatomotor network, dorsal attention network, ventral attention network (VAN), limbic network, FPN, and DMN (**Fig.1D**). The functional connectome was generated by calculating Pearson correlation coefficients between the time series of every possible pair of regions, followed by *r*-to-*z* transformation to enhance data normality. The connectome metrics underwent row-wise thresholding, retaining only the top 10% of connections to ensure that gradient estimation was based on robust connections while minimizing the influence of noisy signals (Dong et al., 2021) . Normalized angular similarity was computed between each pair of rows to construct an affinity matrix, which represents the similarity of functional connectivity profiles across different brain regions (**Fig.1E**). Building upon the affinity matrix, we implemented diffusion map embedding - a nonlinear dimensionality reduction approach - to derive gradient components that effectively characterize the variance patterns within functional connectivity profiles (Coifman & Lafon, 2006). Here, we utilized the conventional manifold learning parameter setting (α = 0.5), in accordance with established gradient mapping literature (Larivière et al., 2020; Margulies et al., 2016), where α modulates the relative contribution of sampling point density to the intrinsic geometry of the underlying manifold. To enable cross-group comparisons of functional gradients, we created a group-level gradient template for two movie conditions, respectively, by using the average functional connectome across all participants in the study cohort with the above-mentioned procedure. Individual gradients were spatially aligned to the group template using Procrustes rotation. The above-mentioned gradient analysis was conducted by using the MATLAB-based Brainspace toolbox (Vos de Wael et al., 2020). Following previous works with gradient analysis, we focused our subsequent analyses on the first three principal gradients, as these typically account for the majority of variance in functional connectivity patterns(**Fig.1F&G**) (Bethlehem et al., 2020; J. Li et al., 2024).

**Figure 1.**
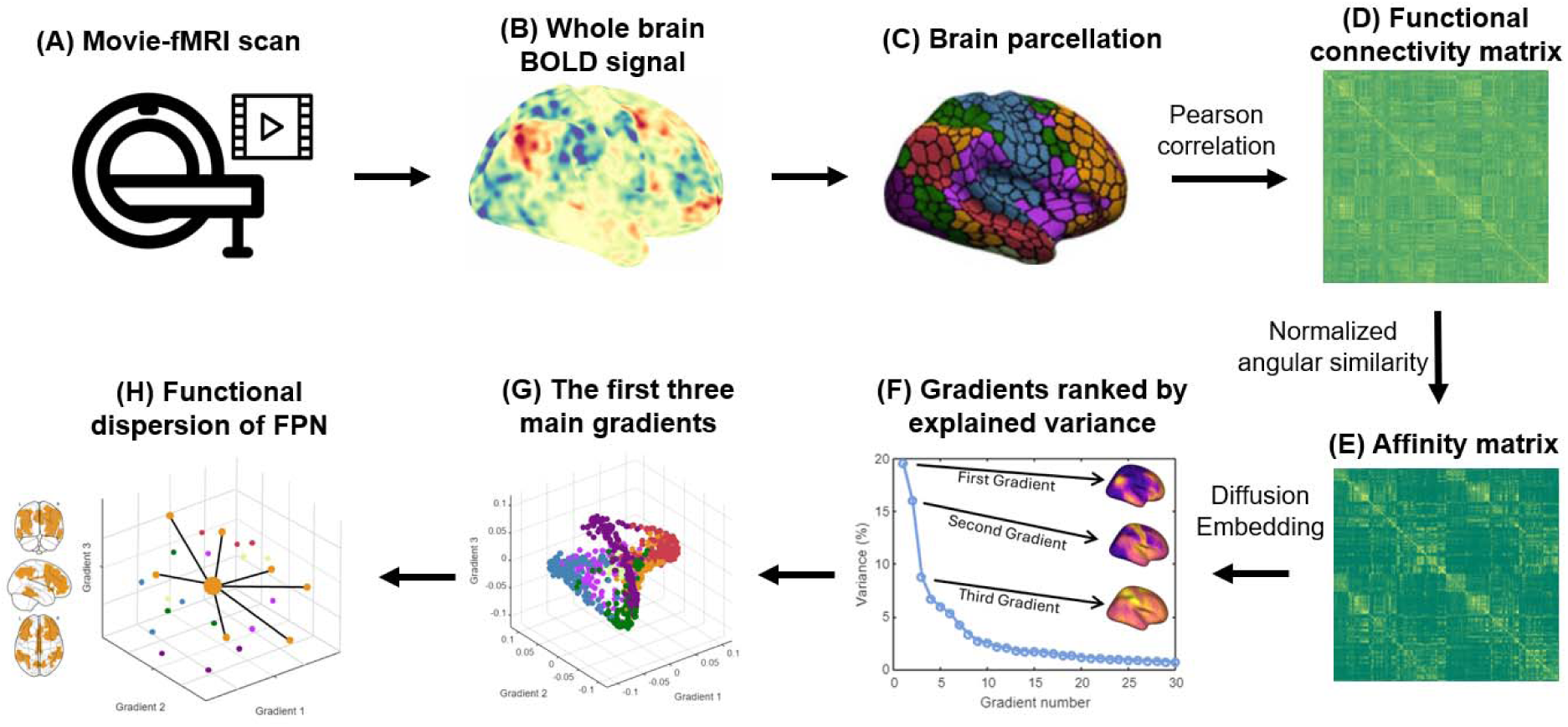
Workflow of FPN dispersion calculation. **(A&B)** Brain activity was recorded while participants watched neutral and negative movie clips in a 7T MRI scanner. **(C&D)** By parcellating the whole-brain cortex into 1,000 regions, the BOLD time series for each region was extracted and pairwise correlated to generate a functional connectivity matrix. **(E)** The affinity matrix was built to represent the similarity of connectivity profiles between brain regions. **(F)** The affinity matrix was then decomposed into a set of principal eigenvectors that explained the most variance using the diffusion embedding approach. **(G)** The first three principal gradients were selected to form a 3-D gradient manifold. **(H)** The functional dispersion of FPN was calculated as the average Euclidean distance between each subregion and network centroid. FPN= frontoparietal network.

### 2.7 FPN functional dedifferentiation

Based on our hypothesis, this study mainly focused on the FPN and its functional dedifferentiation. The functional dispersion of FPN was calculated as the sum of squared Euclidean distances between FPN subregions’ gradient coordinates (i.e., gradient score in the three main gradients) and the network centroids of FPN in the gradient space (**Fig.1H**). The FPN dispersion quantifies the extent of FPN functional dedifferentiation, with higher values reflecting large heterogeneity in network-specific connectivity profiles among FPN regions (i.e., decreased specialization). Although this study primarily focuses on the FPN due to its essential role in emotion regulation, existing evidence suggests that the VAN (also known as the salience network) and DMN are also involved in emotion processing and closely associated with psychiatric disorders (Coutinho et al., 2016; Menon, 2011). Therefore, additional analyses examining VAN and DMN dispersion were conducted, and the results are provided in **Fig.S1&S2**. Notably, we didn’t find any significant correlation between interest behaviors (i.e., emotion regulation difficulties, depression and anxiety symptoms) and the dispersion of VAN or DMN.

In addition to FPN dispersion, we computed the participation coefficient (PC) of the FPN—a graph-theoretical measure widely used to evaluate network dedifferentiation. PC quantifies the extent to which a node’s connections are distributed across distinct functional networks, providing a conventional metric based on functional connectivity (Pedersen M. et al., 2020). A larger PC of the given network indicates that the nodes within the network are more involved in other networks, suggesting increased dedifferentiation. Details of its calculations can be found in the **Supplementary Method**. Here, we included PC to demonstrate that, compared to such traditional functional connectivity metrics (PC), gradient-based measures (i.e., gradient dispersion) may offer more valuable insights by describing network functional architecture from a continuous, multi-dimensional, and macro-scale perspective.

### 2.8 Statistical analyses

We first estimated the associations between emotion regulation difficulty and anxiety/depression symptoms, as well as between FPN dispersion and mental health measures (i.e., DERS score, depression, and anxiety scores) within two age groups. Moreover, linear regression models were constructed to examine the moderating effect of age on the dispersion-behavior relationship by incorporating the age-dispersion interaction term. This procedure was performed with *lm* function in R version 4.4.1.

Mediation analysis was conducted under the premise that FPN dispersion, emotion regulation difficulties, and depression/anxiety symptoms are intercorrelated. This analysis aimed to determine whether emotion regulation mediates the relationship between FPN dispersion and affective disorder symptoms. In the mediation model, the FPN dispersion was set as the independent variable, DERS score as the mediator, and depression/anxiety score as the dependent variable. Mediation analyses were performed using the PROCESS in SPSS (version 30.0, IBM) (Hayes, 2012). The indirect effect of FPN dispersion on depression/anxiety symptoms via emotion regulation difficulty was examined through Model 4 (simple mediation). To address non-normality in the sampling distribution, bias-corrected bootstrapping with 5,000 resamples was applied, and 95% confidence intervals were computed. Thus, a 95% CI excluding zero would indicate statistically significant mediation.

Lastly, to examine how variations in emotional well-being characteristic influence brain organization, we applied latent profile analysis (LPA), a data-driven approach for identifying unobserved subgroups (latent profiles) within a population based on patterns in observed behaviors(Peugh & Fan, 2013; Williams & Kibowski, 2016). Responses to self-report questionnaires (i.e., DERS score, depression score, and anxiety score) were used as indicators to classify individuals within age groups into distinct latent subgroups. These subgroups reflect heterogeneous mental health profiles (e.g., resilient versus vulnerable individuals). By comparing FPN dispersion across the identified subgroups, we assessed whether this neural metric, i.e., FPN dispersion, can effectively distinguish between individuals with distinct mental health profiles. The LPA was conducted with *mclust* package in R version 4.4.1 (Fraley et al., 2012). We specified a range of 1 to 5 potential latent subgroups to allow data-driven identification of the optimal class solution(Spurk et al., 2020). Model selection was based on the Bayesian Information Criterion (BIC), with the best-fitting model determined by the lowest BIC value after evaluating all possible covariance of 14 structures across cluster configurations (for more information see Morgan et al., 2016; Tein et al., 2013).

Age, sex, total intracranial volume (TIV), and mean FD were incorporated into our analyses as additional covariates. The TIV of each subject was estimated after the brain extraction from the T1 image using the Computational Anatomy Toolbox (CAT12) (Gaser et al., 2024).

## 3. Results

### 3.1 Difficulties in emotion regulation are similarly associated with symptoms of depression and anxiety in both younger and older adults

The demographic information and questionnaire responses of the participants were summarized in **Tab.1**. Compared to younger adults, older adults demonstrated lower levels of emotion regulation difficulty (*t* = 6.429, *p* < 0.001, *d* = 1.080), and reported fewer symptoms of depression (*t* = 6.929, *p* < 0.001, *d* = 1.156) and anxiety (*t* = 8.562, *p* < 0.001, *d* = 1.441). Regarding emotional responses to movie clips, older adults exhibited stronger negative emotional valence following exposure to the negative movie (*t* = 2.006, *p* =0.048, *d* = 0.340), although the two groups showed no significant difference across other emotion ratings.

Group-wise correlation analyses revealed significant positive correlations between emotion regulation difficulty and depression/anxiety symptoms within each age group (**Fig.2**). Linear regression results further indicated that age did not moderate the relationship between emotion regulation difficulty and anxiety/depression symptoms, thus suggesting that the positive correlation between emotion regulation difficulty and depression anxiety symptoms are not influenced by age.

**Figure 2.**
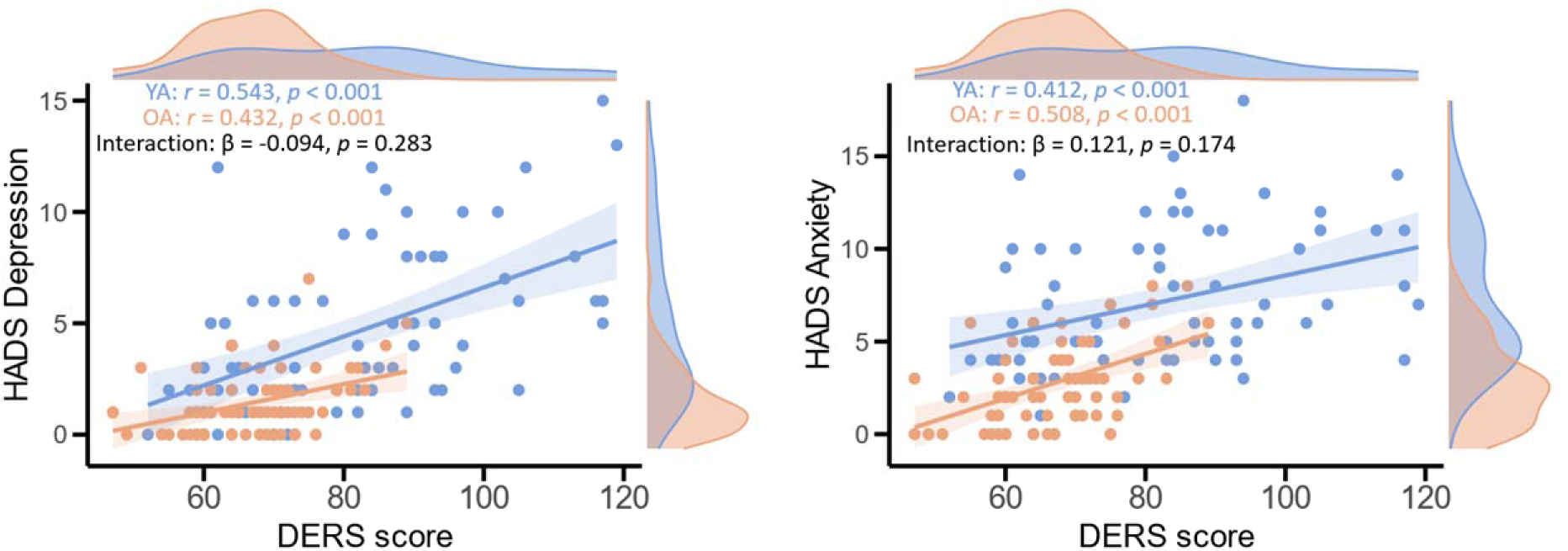
Association between emotion regulation difficulty and depression/anxiety symptoms across two age groups. The correlation analyses revealed a robust positive correlation between emotion regulation difficulty and depression/anxiety symptoms across both age groups. HADS= Hospital Anxiety and Depression Scale; DERS= Difficulties in Emotion Regulation; YA=Younger adults; OA= Older adults.

### 3.2 FPN dispersion is differentially associated with mental health measures in younger than older adults

The FPN dispersion for each movie condition was calculated based on the 3-D gradient space defined by the first three main gradients. These three principal gradients explained 44.39% and 37.39% of the variance in the functional connectivity profiles under negative and neutral conditions, respectively (**Fig.S3**). The details of the main gradients are presented in **Fig.S4**.

Older adults exhibited significantly greater FPN dispersion compared to younger adults under both movie conditions (neutral movie: younger adults = 0.055 ± 0.008,older adults= 0.060 ± 0.010, *t* = -3.179, *p* = 0.002, *d* = -0.540; negative movie: younger adults = 0.059 ± 0.011,older adults= 0.070 ± 0.012, *t* = -5.417, *p* < 0.001, *d* = -0.681), suggesting enhanced FPN functional dedifferentiation in older adults.

Group-wise correlation analyses were conducted separately within each age group to examine the association between FPN dispersion and mental health measures. Under the negative movie condition, FPN dispersion was negatively correlated with DERS scores in younger adults (*r* = -0.254, *p* = 0.044), whereas a positive correlation was observed in older adults (*r* = 0.252, *p* = 0.037), with linear regression model with age-dispersion interaction indicating a significant moderating effect of age (β = 0.194, *p* = 0.012, **Fig.3A**). Moreover, FPN dispersion was positively correlated with depression scores in older adults (*r* = 0.263, *p* =0.036) but not in younger adults (*r* = -0.014, *p* = 0.912), and no significant moderating effect of age was found on this relationship (β = -0.094, *p* = 0.283, **Fig.3B**). No significant correlation was found between FPN dispersion and anxiety scores within each age group (Younger adults: *r* = -0.223, *p* = 0.068; Older adults: *r* = 0.164, *p* =0.196), yet a significant moderating effect of age was revealed (β = 0.174, *p* = 0.014, **Fig.3C**). No significant associations and modulation effects were observed under neutral movie conditions (**Fig.S5**).

**Figure 3.**
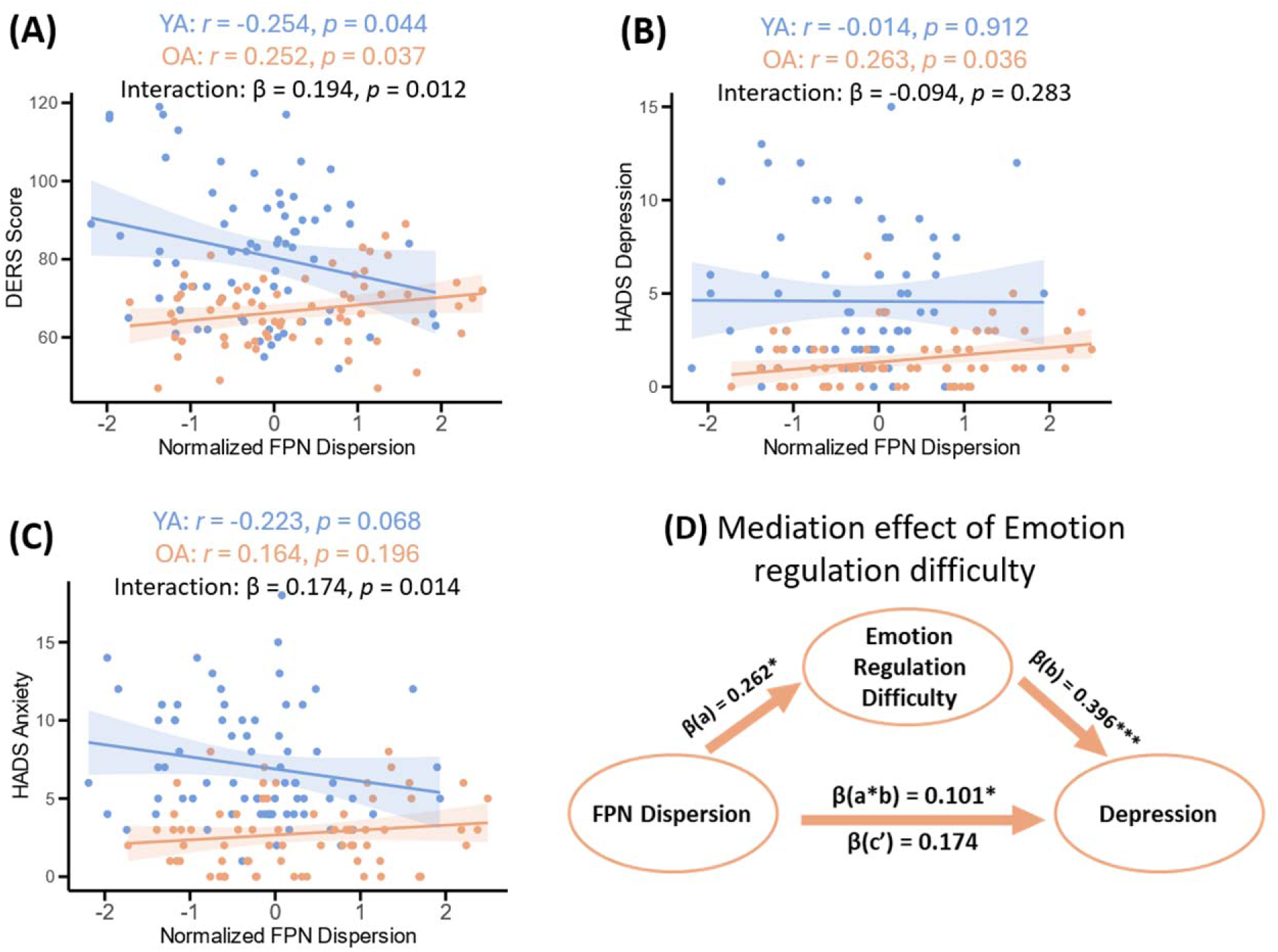
Relationship between FPN dispersion during negative movie watching, emotion regulation difficulty, and depression/anxiety symptoms. **(A)** The scatter plot presents the association between FPN dispersion and emotion regulation difficulty. **(B)** The scatter plot presents the association between FPN dispersion and depressive symptoms. **(C)** The scatter plot presents the association between FPN dispersion and anxiety symptoms. **(D)** Emotion regulation difficulty fully mediated the association between FPN dispersion and depressive symptoms in older adults. HADS= Hospital Anxiety and Depression Scale; DERS= Difficulties in Emotion Regulation; YA=Younger adults; OA= Older adults. FPN= frontoparietal network. *: p <0.05; ***: p<0.001.

Moreover, we computed the participation coefficient of FPN and correlated it with mental health measures. Although we found that younger adults showed lower FPN functional connectivity strength than older people in both two movie conditions (neutral movie: younger adults = 0.835 ± 0.030, older adults = 0.848 ± 0.040, *t* = -2.079, *p* = 0.040, *d* = -0.354; negative movie: younger adults = 0.748 ± 0.033, older adults = 0.766 ± 0.061, *t* = -2.122, *p* = 0.036, *d* = -0.364), no significant correlation was found between FPN functional connectivity strength and mental measures across age groups and movie conditions (all *P*’s > 0.05, **Fig.S6**).

### 3.3 Emotion regulation difficulty mediates the FPN dispersion and depression association

Given that significant associations were exclusively observed among FPN dispersion, DERS score, and depression score within older adults group, we constructed a mediation model examining these relationships. The mediation analysis revealed a significant indirect effect of FPN dispersion through DERS score on depression score (indirect effect = 0.101, 95%CI = [0.004,0.230]). Additionally, the direct effect of FPN on dispersion on depression score was not significant after controlling for DERS score (direct effect = 0.174, *p* = 0.165), indicating full mediation effect of DERS score (**Fig.3D**). These findings suggest that among older adults, increased FPN dispersion (i.e., greater FPN functional dedifferentiation) contributes to depressive symptoms predominantly through impaired emotion regulation, rather than via the direct pathway.

### 3.4 Differences of FPN dispersion in subgroups with diverse mental health profiles

Given substantial evidence supporting the existence of distinct subgroups with varying mental health profiles ranging from minimal to pronounced symptoms (L. Cui et al., 2023; Salami et al., 2018), we next conducted LPA to identify subgroups that likely reflect individuals with normative versus at-risk mental health profiles, using all three measures of emotional regulation, depression and anxiety simultaneously.

For each age group, LPA identified two subgroups. The details of the optimal model selection are presented in **Fig.S7&S8**. In the 2-profile model solution of younger adults, 48 (67%) participants were classified to profile 1 as the resilient group, and 24 (33%) participants to profile 2 as the vulnerable group (**Fig.4A**). No age difference was observed between two subgroups (*t* = -1.192, *p* = 0.241, *d* = -0.325). The resilient group exhibited significantly lower scores compared to the vulnerable group on the difficulty in emotion regulation questionnaire (Resilient group = 73.90 ± 1.73, Vulnerable group = 98.00 ± 3.10, *t* = - 6.275, *p* < 0.001, *d* = -1.702), depression (Resilient group = 2.69 ± 0.24, Vulnerable group = 8.60 ± 0.69, *t* = -8.608, *p* < 0.001, *d* = -2.615), and anxiety (Resilient group = 5.47 ± 0.34, Vulnerable group = 10.52 ± 0.74, *t* = -6.535, *p* < 0.001, *d* = -1.863). However, no significant differences in FPN dispersion were observed between resilient and vulnerable groups among younger adults (Neutral movie: Resilient group = 0.059 ± 0.010, Vulnerable group = 0.060 ± 0.011, *t* = -0.183, *p* = 0. 856, *d* = -0.049; Negative movie: Resilient group = 0.056 ± 0.009, Vulnerable group = 0.054 ± 0.010, *t* = 0.948, *p* = 0.348, *d* = 0.249, **Fig.4B&C**).

**Figure 4.**
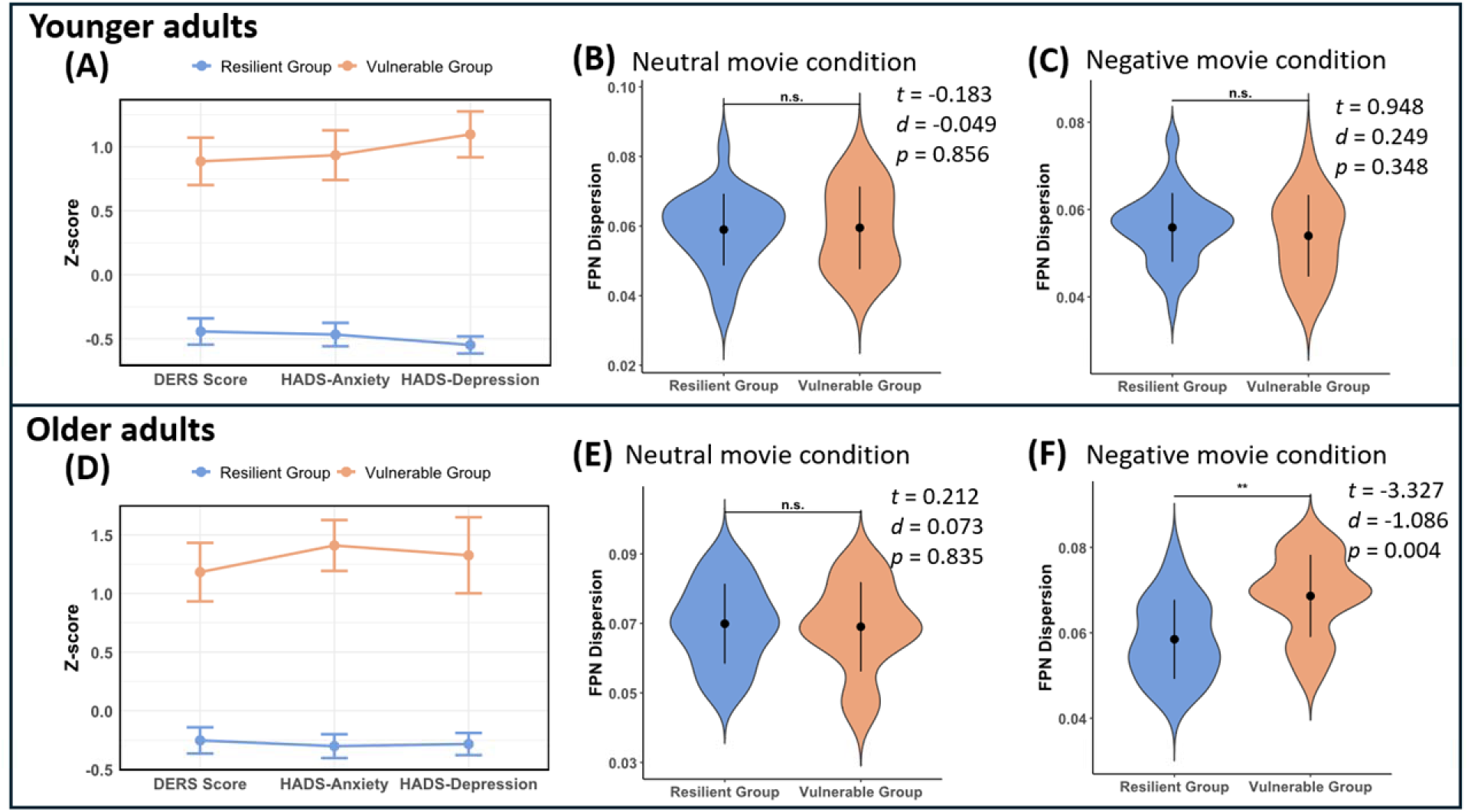
FPN dispersion in two subgroups within each age group. The subgroups were identified based on mental health profiles and their differences in FPN dispersion. **(A)** Latent profile analysis identified two distinct subgroups with different mental health profiles in younger adults. **(B&C)** No significant differences in FPN dispersion between resilient and vulnerable groups were observed in younger adults for either movie condition. **(D)** Latent profile analysis identified two distinct subgroups with different mental health profiles in older adults. **(E)** No significant differences in FPN dispersion between resilient and vulnerable groups were observed in older adults for neutral movie conditions. **(F)** Resilient group showed significant lower FPN dispersion during negative movie watching than vulnerable groups in older adults. HADS= Hospital Anxiety and Depression Scale; DERS= Difficulties in Emotion Regulation; FPN= frontoparietal network. **: *p*<0.01; n.s.= not significant.

In the 2-profile model solution of older adults, 56 (82%) participants were classified to profile 1 as resilient group, and 12 (18%) participants to profile 2 as vulnerable group (**Fig.4D**). No age difference was observed between two subgroups (*t* = -0.302, *p* = 0.767, *d* = -0.111). The RG exhibited significantly lower scores compared to the vulnerable group on DERS (RG = 64.12 ± 2.33, Vulnerable group = 85.30 ± 3.45, *t* = -4.770, *p* < 0.001, d = -1.033), depression (RG = 1.02 ± 0.14, Vulnerable group = 3.33 ± 0.47, *t* = - 4.770, *p* < 0.001, *d* = -2.033), and anxiety (RG = 2.09 ± 0.22, Vulnerable group = 5.83 ± 0.47, *t* = -4.770, *p* < 0.001, *d* = -2.253). Moreover, further analysis indicated lower FPN dispersion under negative movie condition in RG compared to VG (RG = 0.058 ± 0.011, Vulnerable group = 0.069 ± 0.011, *t* = -3.327, *p* = 0.004, *d* = -1.086, **Fig.4F**). No significant results were revealed in neutral movie condition (RG = 0.070 ± 0.012, Vulnerable group = 0.069 ± 0.012, *t* = 0.212, *p* = 0.834, *d* = 0.073, **Fig.4E**). Given the small sample size of the vulnerable group within each age group, additional statistical analyses were performed to verify the robustness of the present findings (Winter, 2013). Results confirmed the reliability of our main findings by showing that the significant difference in FPN dispersion remains between resilient and vulnerable group in older adults after applying Welch’s *t*-test and Mann–Whitney *U* test (**Tab.S1**).

Group-wise LPA revealed that FPN dispersion differentiated only between resilient and vulnerable subgroups in older adults. To examine whether this effect generalized beyond age-specific groups, we conducted whole-sample LPA using age-adjusted mental health profiles and compared FPN dispersion across the resulting subgroups. Three subgroups were identified in LPA based on the optimal model fitting with the highest BIC of -1103.73 (**see Fig.S9A**), including the vulnerable group (N=31), the moderate risk group(N=83), and the resilient group (N=26) (**Fig.S9B)**. However, FPN dispersion showed no significant differences across subgroups in either neutral (*F* (2,137) = 0.971, *p* = 0.381, η² = 0.014) or negative (*F* (2,137) = 0.162, *p =* 0.851, η² = 0.002) movie conditions (**Fig.S9C&D**). The findings further underscored the distinctive relationship, which is specific to older adults, between FPN dispersion and mental health profiles.

## 4. Discussion

Developing validated neuroimaging biomarkers that can identify predispositions to late-life mood disorders is critical for implementing targeted preventive interventions in geriatric mental healthcare (Hannon & Bijsterbosch, 2023). Using an ecologically valid, naturalistic fMRI paradigm combined with a multidimensional gradient framework, we assessed the functional dispersion of the FPN to quantify its functional organization during real-life emotional experiences. We demonstrated that older adults exhibited greater FPN dedifferentiation during naturalistic processing, reflected by increased FPN dispersion. Older adults with increased FPN dispersion during negative movie watching reported higher levels of affective disorder symptoms, with this relationship being fully mediated by diminished emotion regulation capacity. Importantly, older adults with vulnerable health profiles exhibited greater FPN dispersion compared to normal resilient age-match groups. These findings suggest that excessive FPN functional dedifferentiation during negative emotional processing is associated with poorer mental health outcomes in older adults and indicate that increased FPN dispersion could serve as a potential neuroimaging biomarker of heightened vulnerability to affective disorders in this population.

Functional dedifferentiation, characterized by reduced neural specificity, has been consistently established in aging populations through mounting fMRI evidence employing diverse methodological approaches (Koen et al., 2020; Koen & Rugg, 2019; Ziaei et al., 2016). For example, older adults have exhibited less specialized responses of the ventral visual cortex during visual recognition (D. C. Park et al., 2004), as well as reduced system segregation of intrinsic functional brain networks in aging population (Chan et al., 2014). Consistent with these findings, we provide novel evidence demonstrating age-related functional dedifferentiation within the FPN during naturalistic movie watching, specifically during negatively valanced stimuli. This observation broadens current neurobiological models of aging by revealing that dedifferentiation manifestation is not only in highly constrained tasks or during the resting state, but also observable during dynamic real-world like stimuli. The increased FPN dispersion reflects diminished similarity within the FPN across multidimensional spaces of functional connectivity profiles, indicating an age-related decline in the network’s capacity to maintain functional specialization during naturalistic processing (Redcay & Moraczewski, 2020).

Moreover, FPN dispersion was negatively correlated with mental health measures in older adults, whereas their younger counterparts exhibited either no association or the opposite pattern. Previous studies have suggested that age-related neural degradation can lead to reduced neural efficiency, manifesting as a lower signal-to-noise ratio and slower neural transmission speed (S.-C. Li et al., 2001; S.-C. Li & Rieckmann, 2014). Critically, younger brains appear to maintain an optimal specificity-integration balance in functional network topology through high neural efficiency, which allows transient shifts toward integrated processing under cognitive demands without compromising specialized computations (Ezaki et al., 2018). Conversely, aging brains exhibit a maladaptive tipping of this topological balance, whereby age-associated neural degradation makes older adults more vulnerable to functional dedifferentiation (Cabral et al., 2017; Uddin, 2021; Yin et al., 2016). Previous studies have demonstrated that functional dedifferentiation is a strong predictor of age-related cognitive decline, particularly affecting higher-order cognitive functions such as working memory and reasoning ability (Bethlehem et al., 2020; Korkki et al., 2025; R. Pedersen et al., 2021, 2023). Likewise, during emotional processing, FPN dedifferentiation in older adults may disrupt specialized neural processing essential for adaptive emotional functioning, such as inhibitory control over automatic emotional responses, flexible prefrontal regulation and context-related allocation of attentional resources (Dalgleish, 2004; Fossati, 2012). Our mediation analysis aligns with the idea that increased FPN dispersion among older adults contributes to depressive symptomatology through impaired emotion regulation capacity. Collectively, these findings suggest that younger brains may reduce specialized FPN for processing cognitive-emotional functions without compromising the functional network structure, whereas older brains undergo pronounced dedifferentiation that disrupts both functional segregation and adaptive integration, contributing to impaired emotion regulation capacity and elevated depressive symptoms.

Our latent profile analysis identified distinct patterns of FPN dispersion that differentiated resilient from vulnerable subgroups among older adults, suggesting that FPN functional architecture may serve as a neural signature of affective vulnerability in aging. Specifically, resilient older adults appear capable of maintaining an optimal balance in the network organization (i.e., exhibiting relatively low dedifferentiation) that supports emotional adaptation without compromising functional network boundaries. In contrast, vulnerable individuals exhibit pronounced dedifferentiation, which destabilizes the FPN’s dual functionality in cognitive control and emotion regulation (W. Li et al., 2021; Ray et al., 2020). A recent study further supports this view, demonstrating increased FPN-related inter-network connectivity in participants experiencing catastrophic events, suggesting that FPN dedifferentiation is associated with post-stress vulnerability (Mas-Cuesta et al., 2024). Given that our additional analyses with salience and DMN did not reveal such pattern as for FPN, we suggest that FPN, which is important in mediating top-down affective control and coordinating cognitive-emotional integration, is uniquely sensitive in detecting early disruptions at the cognitive-emotional interaction (Morawetz et al., 2020; Morawetz & Basten, 2024) or dysregulation of emotion. This sensitivity enables the identification of older adults at heightened risk for affective disorders—specifically, individuals who exhibit symptoms yet remain highly vulnerable to disturbances triggered by negative events. Although not directly tested, but one extension of these findings is related to emotional well-being literature where older adults were reported higher levels of positive emotions (Carstensen et al., 2011). With current findings we can speculate that perhaps the FPN organization might play an important role in age-related shifts in processing emotional cues or alteration in brain organization involved in emotional processing.

Notably, the significant results described above were exclusively observed during negative movie viewing, underscoring the selectivity of the FPN and its pivotal role in responding to emotionally charged real-world scenarios. Unlike emotionally neutral contexts where FPN primarily supports generalized cognitive control, exposure to negative stimuli adds extra demands on the network to integrate and regulate emotion, thus, increasing its sensitivity in detecting dysfunction in emotional processing (Gruskin et al., 2020; Kirk et al., 2023). Thus, the superior utility of negative clips compared to neutral clips in examining emotion-related behaviors stems from their heightened ecological validity and capacity to impose dynamic emotional loads (Kirk et al., 2022; Wang et al., 2023).

While this study advances the understanding of FPN dedifferentiation in aging and its relevance to mental health, several limitations warrant consideration. First, the current sample consists of healthy participants, which constrains generalizability to clinical populations with affective disorders. This necessitates future validation in aging cohorts spanning subclinical to patient groups to further validate whether FPN could be a generalized and reliable neural signature for late-life affective disorders. Second, while we identified FPN dedifferentiation as a neural correlate of emotional vulnerability, the underlying neuromodulatory mechanisms—particularly dopamine and norepinephrine systems, known to be critical for network dedifferentiation (Mooraj et al., 2025; Nordin et al., 2022; R. Pedersen et al., 2023)—remain uncharacterized. Multimodal approaches integrating receptor-specific PET imaging and genetic profiling could disentangle molecular drivers of these effects. Third, the exclusive focus on negative emotional contexts leaves an open question about valence specificity, as aging is also associated with altered processing of positive emotions (e.g., the positivity bias in older adults, Carstensen & Mikels, 2005; Kalenzaga et al., 2016). Incorporating positive emotional stimuli would help determine whether dedifferentiation reflects a general network dysregulation or is specific to negatively-valenced emotional processing.

In conclusion, by combining naturalistic movie-fMRI with gradient analysis, the current study demonstrates that enhanced FPN dedifferentiation during negative emotional processing is associated with depressive symptoms in older adults, an effect mediated by impaired emotion regulation capacity. Furthermore, increased FPN dispersion may represent a potential neuroimaging biomarker for identifying individuals at a heightened risk of affective disorders in aging populations. These findings highlight the pivotal role of the FPN in age-related emotional vulnerability, offering valuable insights and potential targets for early detection and preventive interventions aimed at improving mental health in late life.

**Table 1.** Demographic information and studied variables.

|  | Younger Group (N=72) | Older Group (N=68) | t-value |
| --- | --- | --- | --- |
| Age(yrs) | 25.75 ± 4.25 | 70.90 ± 3.83 | - |
| TIV (cm<sup>3</sup>) | 1458.68 ± 207.82 | 1503.52 ± 191.11 | -1.327 |
| Head motion (mm) |  |  |  |
| Neutral movie | 0.13 ± 0.08 | 0.20 ± 0.08 | -5.600*** |
| Negative movie | 0.13 ± 0.06 | 0.21 ± 0.09 | -6.617*** |
| DERS score | 81.60 ± 17.20 | 66.85 ± 8.87 | 6.429*** |
| HADS |  |  |  |
| Depression score | 4.58 ± 3.57 | 1.43 ± 1.44 | 6.929*** |
| Anxiety score | 7.08 ± 3.66 | 2.75 ± 2.19 | 8.562*** |
| EV rating |  |  |  |
| Baseline | 6.07 ± 1.28 | 6.19 ± 1.69 | -0.479 |
| Neutral movie | 6.00 ± 1.31 | 5.91 ± 1.79 | 0.334 |
| Negative movie | 3.81 ± 1.50 | 3.25 ± 1.77 | 2.006* |
| EA rating |  |  |  |
| Baseline | 4.79 ± 1.54 | 4.25 ± 1.88 | 1.860 |
| Neutral movie | 4.65 ± 1.89 | 4.18 ± 2.00 | 1.445 |
| Negative movie | 6.19 ± 1.58 | 6.09 ± 1.79 | 0.373 |

Note: TIV=total intracranial volume; DERS=Difficulties in Emotion Regulation Scale; HADS=Hospital Anxiety and Depression Scale; EV=emotional valence; EA=emotional arousal; \*: $p<0.05$ , \*\*: $p<0.01$ , \*\*\*: $p<0.001$ .

## Acknowledgments

We would like to thank radiographers and MR physicists at the 7T MR center at NTNU for their help during this project. We also would like to thank our participants for their time and effort during the experiment. We thank Jørgen Østmo-Sæter Olsnes, Leona Rahel Bätz, Stian Framvik, Avneesh Jain, Jae Hong, and Karina Tømmerdal for their help during data collection, and thanks Xiaqing Lan for her help in data curation.

## Data Availability

The data will be available from the senior author, M.Z., upon request.

## CRediT authorship contribution statement

**S.Y.** Conceptualization, Data curation, Methodology, Formal analysis, Visualization, Writing – original draft. **A.D.**: Investigation, Formal analysis, Writing – review & editing. **M.Z.**: Conceptualization, Funding acquisition, Supervision, Project administration, Writing – review & editing. **A.S.**: Supervision, Writing – review & editing.

