## Supplemental Information for "Frontoparietal functional dedifferentiation during naturalistic movie watching among older adults at risk of emotional vulnerability"

**This file includes:**

Supplementary Methods

Supplementary Figures S1 to S9

Supplementary Tables S1

**Supplementary Methods**

**Imaging data preprocessing**

Results included in this manuscript come from preprocessing performed using *fMRIPrep* 22.0.2 (Esteban, Markiewicz, et al. (2018); Esteban, Blair, et al. (2018); RRID:SCR_016216), which is based on *Nipype* 1.8.5 (K. Gorgolewski et al. (2011); K. J. Gorgolewski et al. (2018); RRID:SCR_002502).

**Preprocessing of B0 inhomogeneity mappings**

A total of 2 fieldmaps were found available within the input BIDS structure. A B0-nonuniformity map (or fieldmap) was estimated based on two (or more) echo-planar imaging (EPI) references with topup (Andersson, Skare, and Ashburner (2003); FSL 6.0.5.1:57b01774).

**Anatomical data preprocessing**

A total of 1 T1-weighted (T1w) images were found within the input BIDS dataset. The T1-weighted (T1w) image was corrected for intensity non-uniformity (INU) with N4BiasFieldCorrection (Tustison et al. 2010), distributed with ANTs 2.3.3 (Avants et al. 2008, RRID:SCR_004757), and used as T1w-reference throughout the workflow. The T1w-reference was then skull-stripped with a *Nipype* implementation of the antsBrainExtraction.sh workflow (from ANTs), using OASIS30ANTs as target template. Brain tissue segmentation of cerebrospinal fluid (CSF), white-matter (WM) and gray-matter (GM) was performed on the brain-extracted T1w using fast (FSL 6.0.5.1:57b01774, RRID:SCR_002823, Zhang, Brady, and Smith 2001). Brain surfaces were reconstructed using recon-all (FreeSurfer 7.2.0, RRID:SCR_001847, Dale, Fischl, and Sereno 1999), and the brain mask estimated previously was refined with a custom variation of the method to reconcile ANTs-derived and FreeSurfer-derived segmentations of the cortical gray-matter of Mindboggle (RRID:SCR_002438, Klein et al. 2017). Volume-based spatial normalization to one standard space (MNI152NLin2009cAsym) was performed through nonlinear registration with antsRegistration (ANTs 2.3.3), using brain-extracted versions of both T1w reference and the T1w template. The following template was selected for spatial normalization: *ICBM 152 Nonlinear Asymmetrical template version 2009c* [Fonov et al. (2009), RRID:SCR_008796; TemplateFlow ID: MNI152NLin2009cAsym].

**Functional data preprocessing**

For each of the 2 BOLD runs found per subject (across all tasks and sessions), the following preprocessing was performed. First, a reference volume and its skull-stripped version were generated using a custom methodology of *fMRIPrep*. Head-motion parameters with respect to the BOLD reference (transformation matrices, and six corresponding rotation and translation parameters) are estimated before any spatiotemporal filtering using mcflirt (FSL 6.0.5.1:57b01774, Jenkinson et al. 2002). BOLD runs were slice-time corrected to 0.978s (0.5 of slice acquisition range 0s-1.96s) using 3dTshift from AFNI (Cox and Hyde 1997, RRID:SCR_005927). The BOLD time-series (including slice-timing correction when applied) were resampled onto their original, native space by applying the transforms to correct for head-motion. These resampled BOLD time-series will be referred to as *preprocessed BOLD in original space*, or just *preprocessed BOLD*. The BOLD reference was then co-registered to the T1w reference using bbregister (FreeSurfer) which implements boundary-based registration (Greve and Fischl 2009). Co-registration was configured with six degrees of freedom. Several confounding time-series were calculated based on the *preprocessed BOLD*: framewise displacement (FD), DVARS and three region-wise global signals. FD was computed using two formulations following Power (absolute sum of relative motions, Power et al. (2014)) and Jenkinson (relative root mean square displacement between affines, Jenkinson et al. (2002)). FD and DVARS are calculated for each functional run, both using their implementations in *Nipype* (following the definitions by Power et al. 2014). The three global signals are extracted within the CSF, the WM, and the whole-brain masks. Additionally, a set of physiological regressors were extracted to allow for component-based noise correction (*CompCor*, Behzadi et al. 2007). Principal components are estimated after high-pass filtering the *preprocessed BOLD* time-series (using a discrete cosine filter with 128s cut-off) for the two *CompCor* variants: temporal (tCompCor) and anatomical (aCompCor). tCompCor components are then calculated from the top 2% variable voxels within the brain mask. For aCompCor, three probabilistic masks (CSF, WM and combined CSF+WM) are generated in anatomical space. The implementation differs from that of Behzadi et al. in that instead of eroding the masks by 2 pixels on BOLD space, a mask of pixels that likely contain a volume fraction of GM is subtracted from the aCompCor masks. This mask is obtained by dilating a GM mask extracted from the FreeSurfer’s *aseg* segmentation, and it ensures components are not extracted from voxels containing a minimal fraction of GM. Finally, these masks are resampled into BOLD space and binarized by thresholding at 0.99 (as in the original implementation). Components are also calculated separately within the WM and CSF masks. For each CompCor decomposition, the *k* components with the largest singular values are retained, such that the retained components’ time series are sufficient to explain 50 percent of variance across the nuisance mask (CSF, WM, combined, or temporal). The remaining components are dropped from consideration. The head-motion estimates calculated in the correction step were also placed within the corresponding confounds file. The confound time series derived from head motion estimates and global signals were expanded with the inclusion of temporal derivatives and quadratic terms for each (Satterthwaite et al. 2013). Frames that exceeded a threshold of 0.5 mm FD or 1.5 standardized DVARS were annotated as motion outliers. Additional nuisance timeseries are calculated by means of principal components analysis of the signal found within a thin band (*crown*) of voxels around the edge of the brain, as proposed by (Patriat, Reynolds, and Birn 2017). The BOLD time-series were resampled into standard space, generating a *preprocessed BOLD run in MNI152NLin2009cAsym space*. First, a reference volume and its skull-stripped version were generated using a custom methodology of *fMRIPrep*. All resamplings can be performed with *a single interpolation step* by composing all the pertinent transformations (i.e. head-motion transform matrices, susceptibility distortion correction when available, and co-registrations to anatomical and output spaces). Gridded (volumetric) resamplings were performed using antsApplyTransforms (ANTs), configured with Lanczos interpolation to minimize the smoothing effects of other kernels (Lanczos 1964). Non-gridded (surface) resamplings were performed using mri_vol2surf (FreeSurfer).

Many internal operations of *fMRIPrep* use *Nilearn* 0.9.1 (Abraham et al. 2014, RRID:SCR_001362), mostly within the functional processing workflow. For more details of the pipeline, see [the section corresponding to workflows in *fMRIPrep*’s documentation](https://fmriprep.readthedocs.io/en/latest/workflows.html).

Copyright Waiver

The above boilerplate text was automatically generated by fMRIPrep with the express intention that users should copy and paste this text into their manuscripts *unchanged*. It is released under the [CC0](https://creativecommons.org/publicdomain/zero/1.0/) license.

Copyright Waiver

The above methods description text was automatically generated by *XCP* with the express intention that users should copy and paste this text into their manuscripts *unchanged*. It is released under the [CC0](https://creativecommons.org/publicdomain/zero/1.0/) license.

**Calculation of participation coefficient**

Participation coefficient (PC), a graph theoretical metric that measures how one brain regions connect to other brain regions of different networks (Pedersen et al., 2020). The calculation of PC was performed using unthresholded Fisher r-z transformed connectivity matrices. These matrices were subsequently binarized to create undirected and unweighted graphs, considering only positive connectivity values and applying a sparsity threshold of 0.1 (i.e., only the 10% most stronger connections were retained). Based on the pre-defined seven networks of the Schaefer atlas(Schaefer et al., 2018), the participation coefficient for each region (or node) can be calculated as the following formula:

$$PC_{i}=1-\sum_{m=1}^{M} \left( \frac{k_{im}}{k_{i}} \right)^{2}$$

Based on the binary graph, *m* represents a specific module within the set of modules *M*. The term *k_i_* denotes the total number of connections associated with node *i* across the entire graph, while *k_im_* refers to the number of connections between node *i* and module *m*. For each participant, the FPN PC was obtained by averaging across PC of FPN regions to quantify the functional integration of FPN, and a higher FPN PC indicates increased functional dedifferentiation.

**Supplementary Figures**

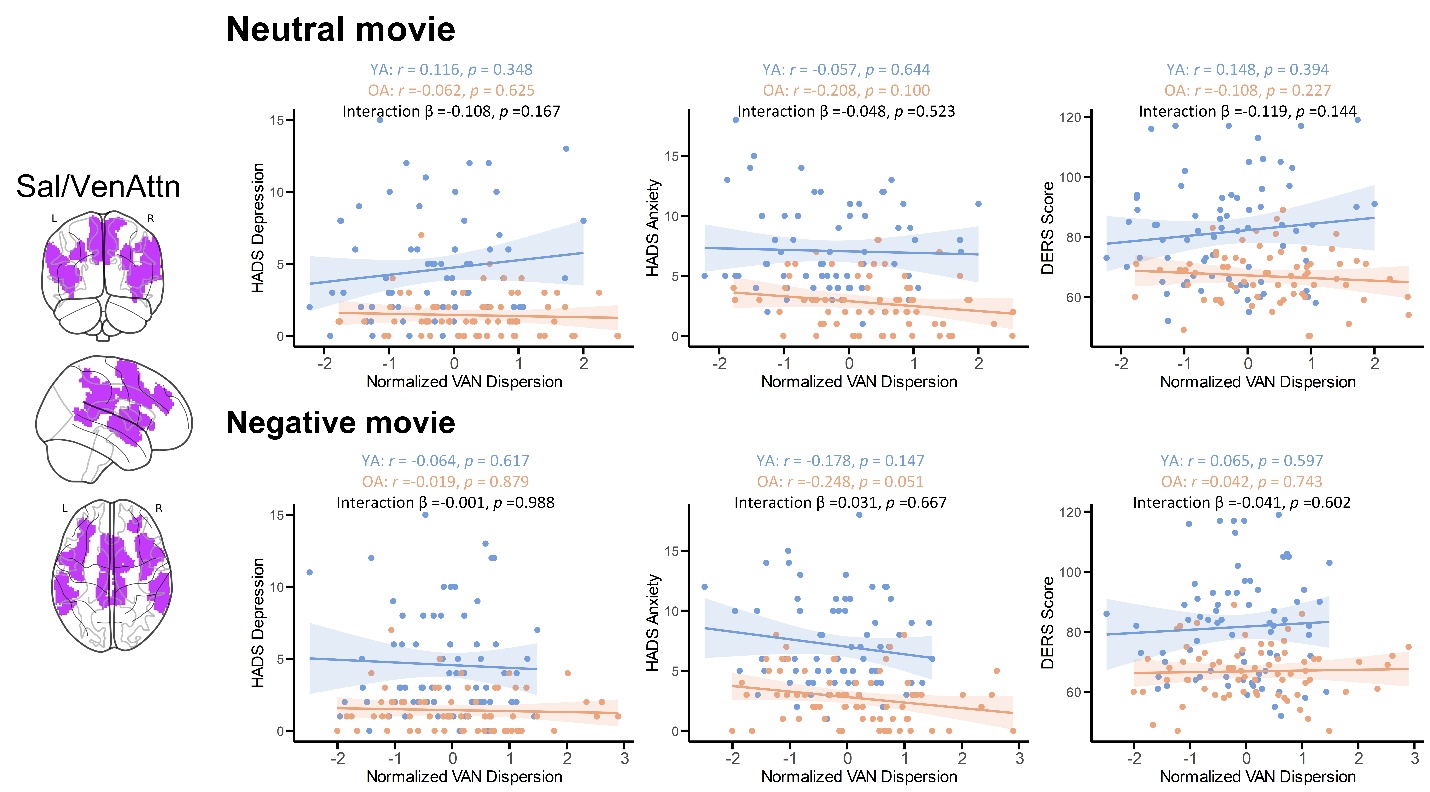

**Figure S1. Relationship between ventral attention network (VAN) dispersion during both neutral and negative movie watching, emotion regulation difficulty, and depression/anxiety symptoms.** HADS= Hospital Anxiety and Depression Scale; DERS= Difficulties in Emotion Regulation; YA=Younger adults; OA= Older adults.

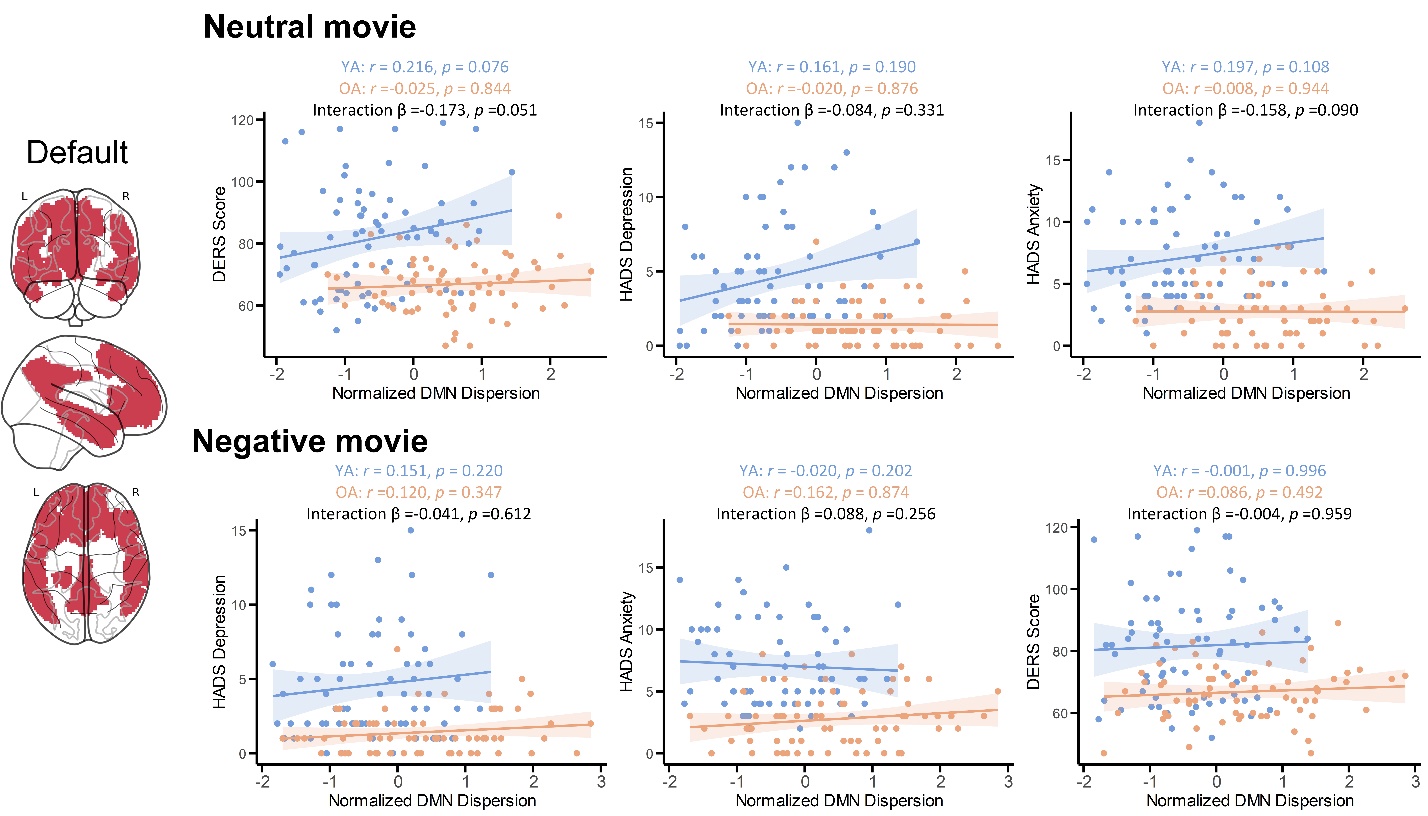

**Figure S2. Relationship between default mode network (DMN) dispersion during both neutral and negative movie watching, emotion regulation difficulty, and depression/anxiety symptoms.** HADS= Hospital Anxiety and Depression Scale; DERS= Difficulties in Emotion Regulation; YA=Younger adults; OA= Older adults.

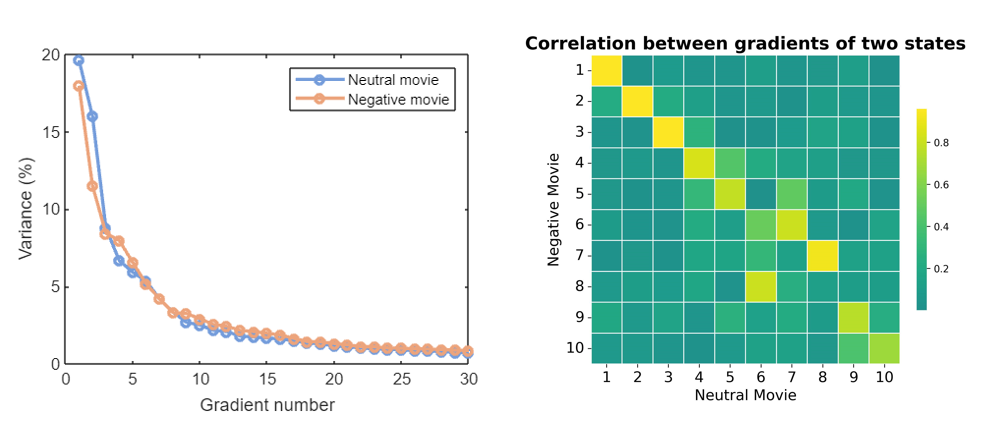

**Figure S3. The averaged explained ratio of the first 30 diffusion embedding components under neutral and negatives movie condition, and the spatial correlations of the averaged gradient maps between two conditions.** The first three principal gradients explained 44.39% and 37.39% of the total variance in the functional connectivity profiles under negative and neutral conditions, respectively. The main gradients are highly correlated between two conditions (Gradient 1: r=0.944, p <0.001; Gradient 2: r=0.942, p <0.001; Gradient 3: r=0.959, p <0.001), indicating a high similarity of the gradient manifolds between two conditions.

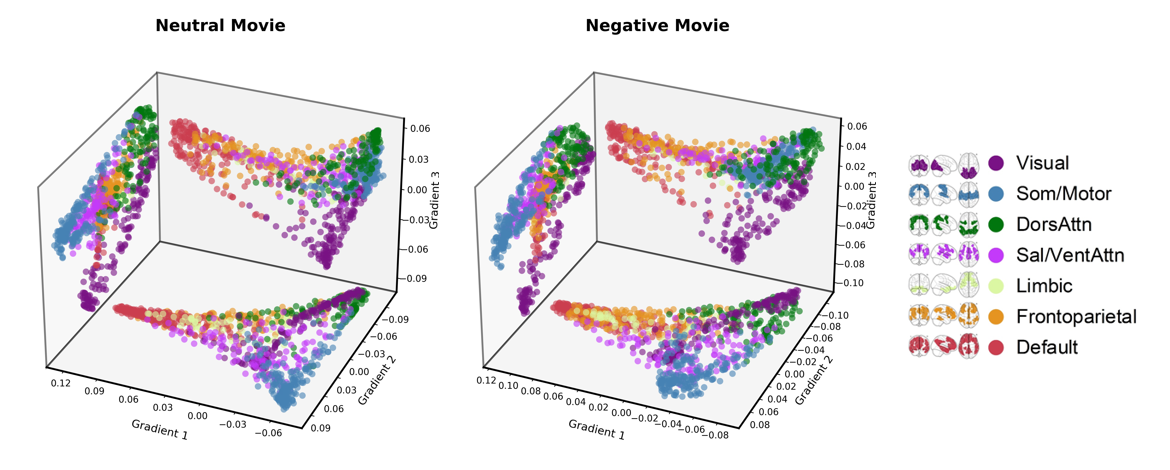

**Figure S4. The first three main gradients of neutral and negative movie condition.** The scatter plots present 2D projections of the gradients in 3D space and are color-coded by parcel assignments of Yeo’s 7 functional networks. The gradient representations are similar between two conditions. The first gradient (gradient 1) spans from unimodal areas, including visual cortex and sensorimotor areas, to association areas, recognized as the sensory-association axis. The second gradient (gradient 2) forms a visual to sensorimotor axis. The third gradient (gradient 3) separates visual regions from other cortical areas, capturing a visual-nonvisual axis.

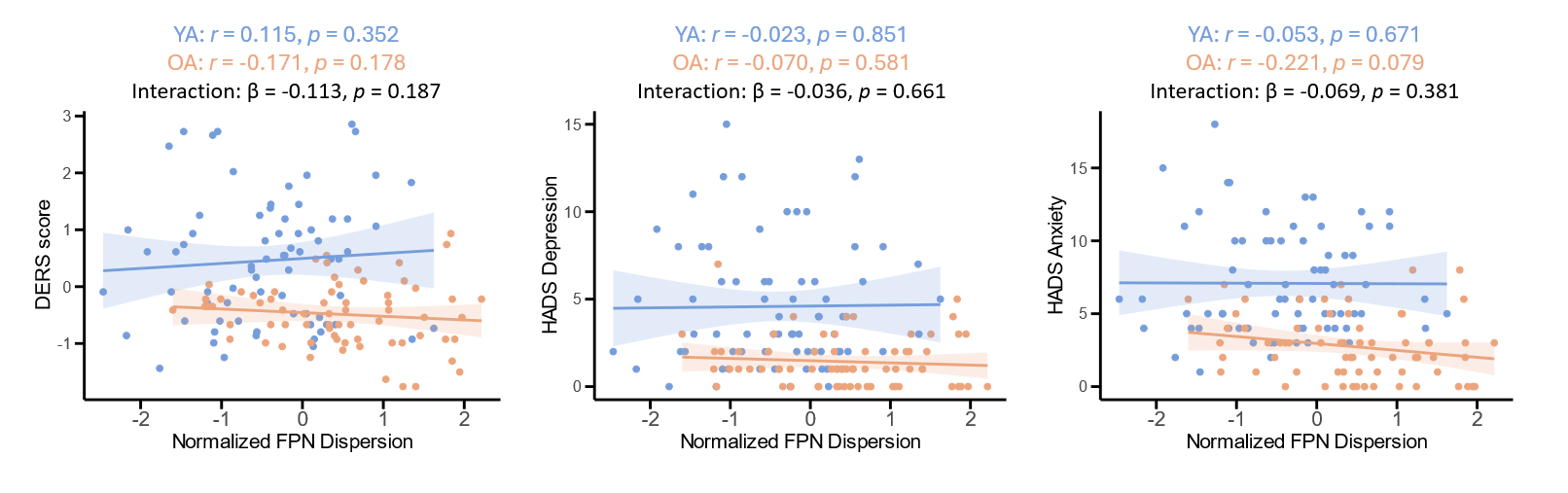

**Figure S5. Relationship between FPN dispersion during neutral movie watching, emotion regulation difficulty, and depression/anxiety symptoms.** HADS= Hospital Anxiety and Depression Scale; DERS= Difficulties in Emotion Regulation; YA=Younger adults; OA= Older adults. FPN= frontoparietal network.

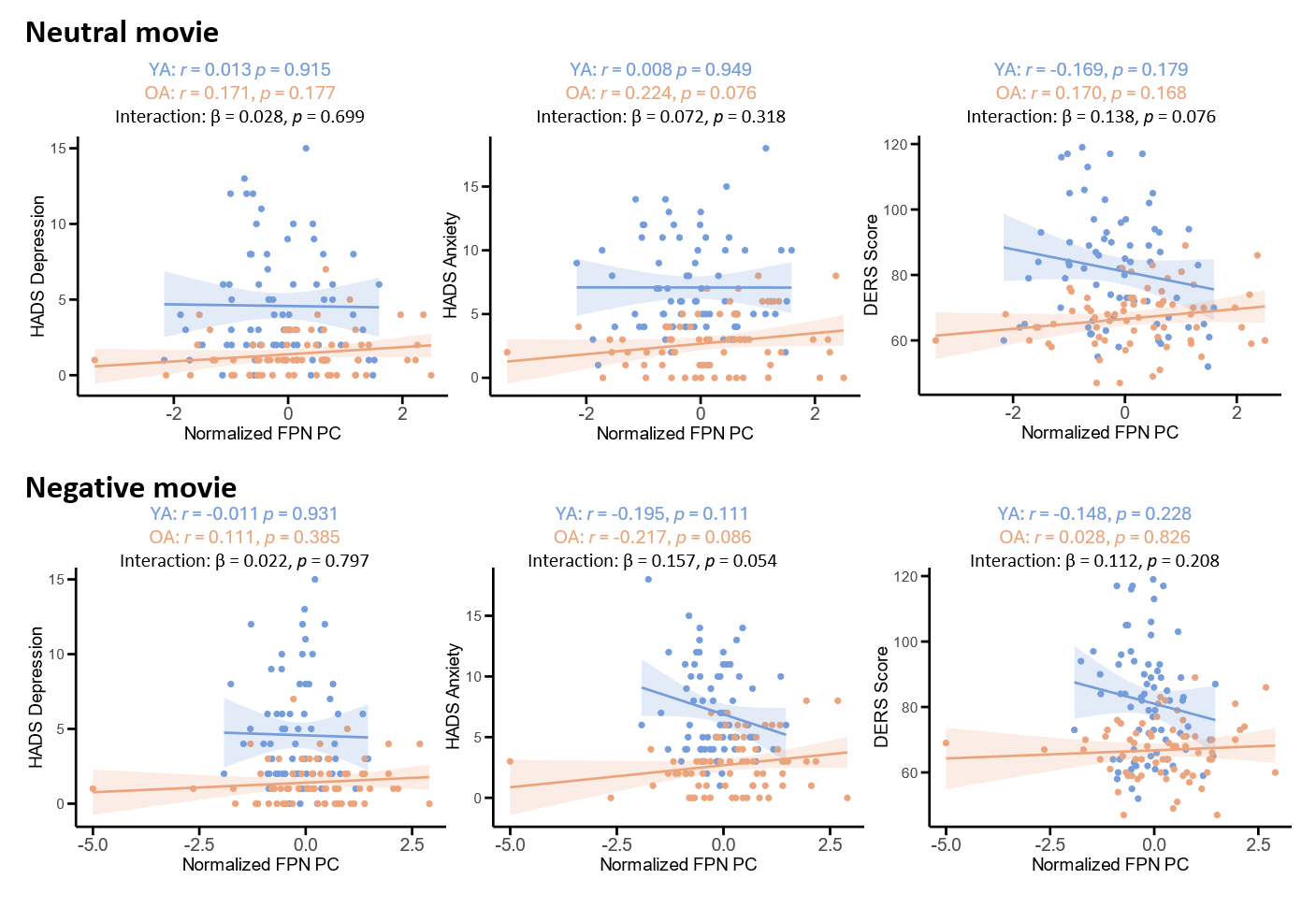

**Figure S6. Relationship between FPN participation coefficient during neutral movie watching, emotion regulation difficulty, and depression/anxiety symptoms.** HADS= Hospital Anxiety and Depression Scale; DERS= Difficulties in Emotion Regulation; YA=Younger adults; OA= Older adults. FPN= frontoparietal network, PC=participation coefficient.

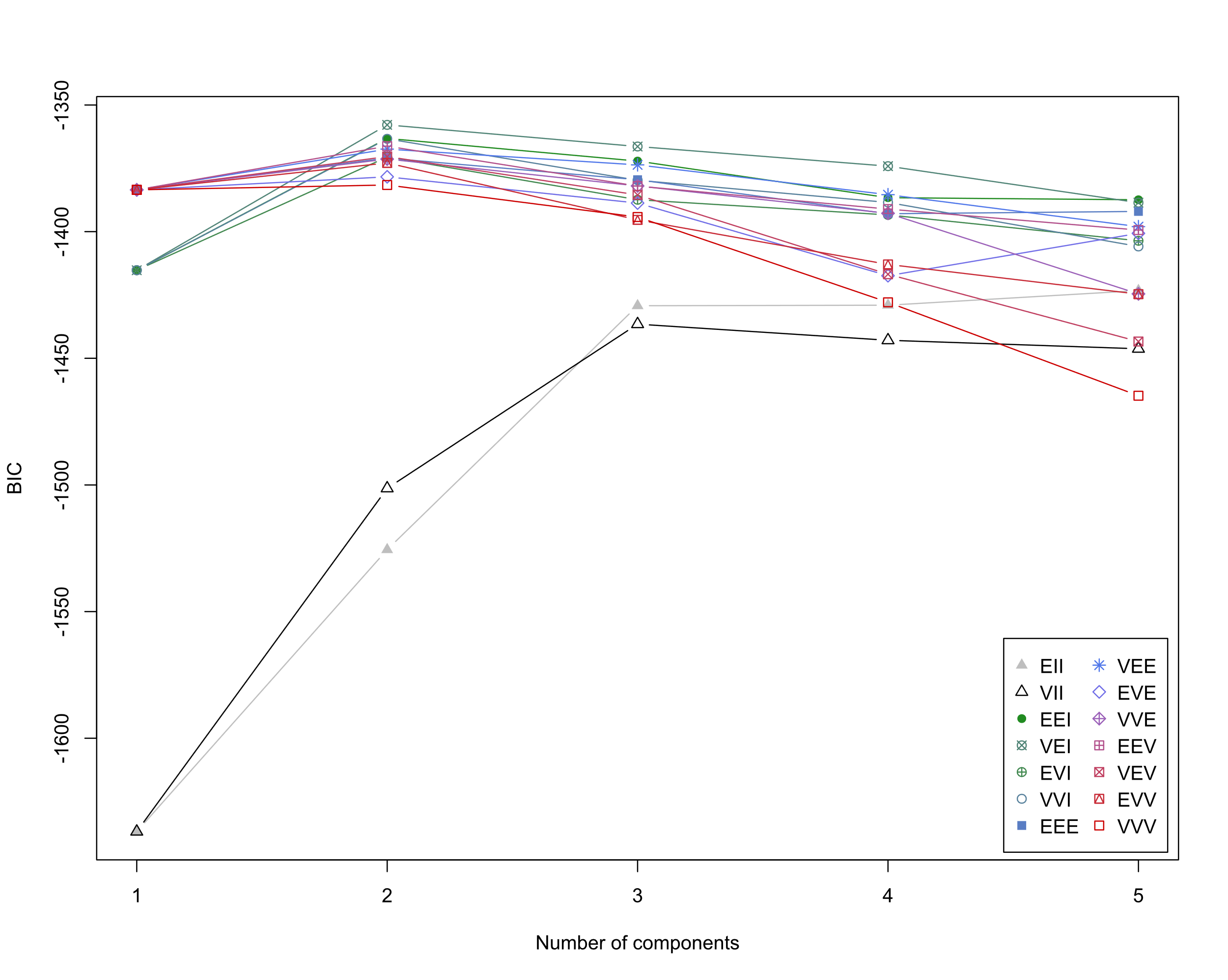

**Figure S7. The Bayesian Information Criterion (BIC) values for different numbers of latent profiles in younger adults.** The best-fitting model was identified as a VEI (diagonal, equal shape) model with 2 mixture components (BIC = -1357.85).

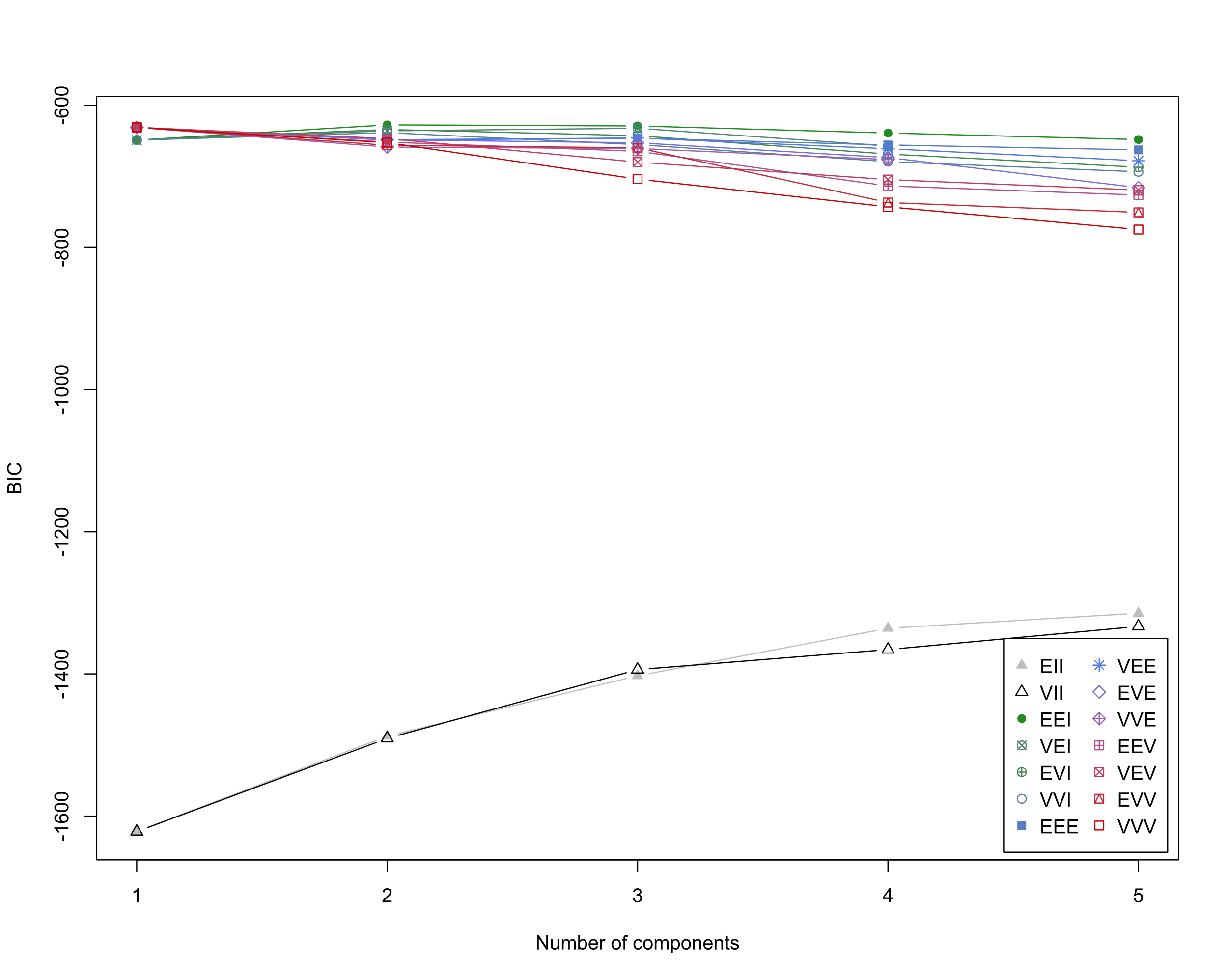

**Figure S8. The Bayesian Information Criterion (BIC) values for different numbers of latent profiles in older adults.** The best-fitting model was identified as a EEI (diagonal, equal volume and shape) model with 2 mixture components (BIC = -617.73).

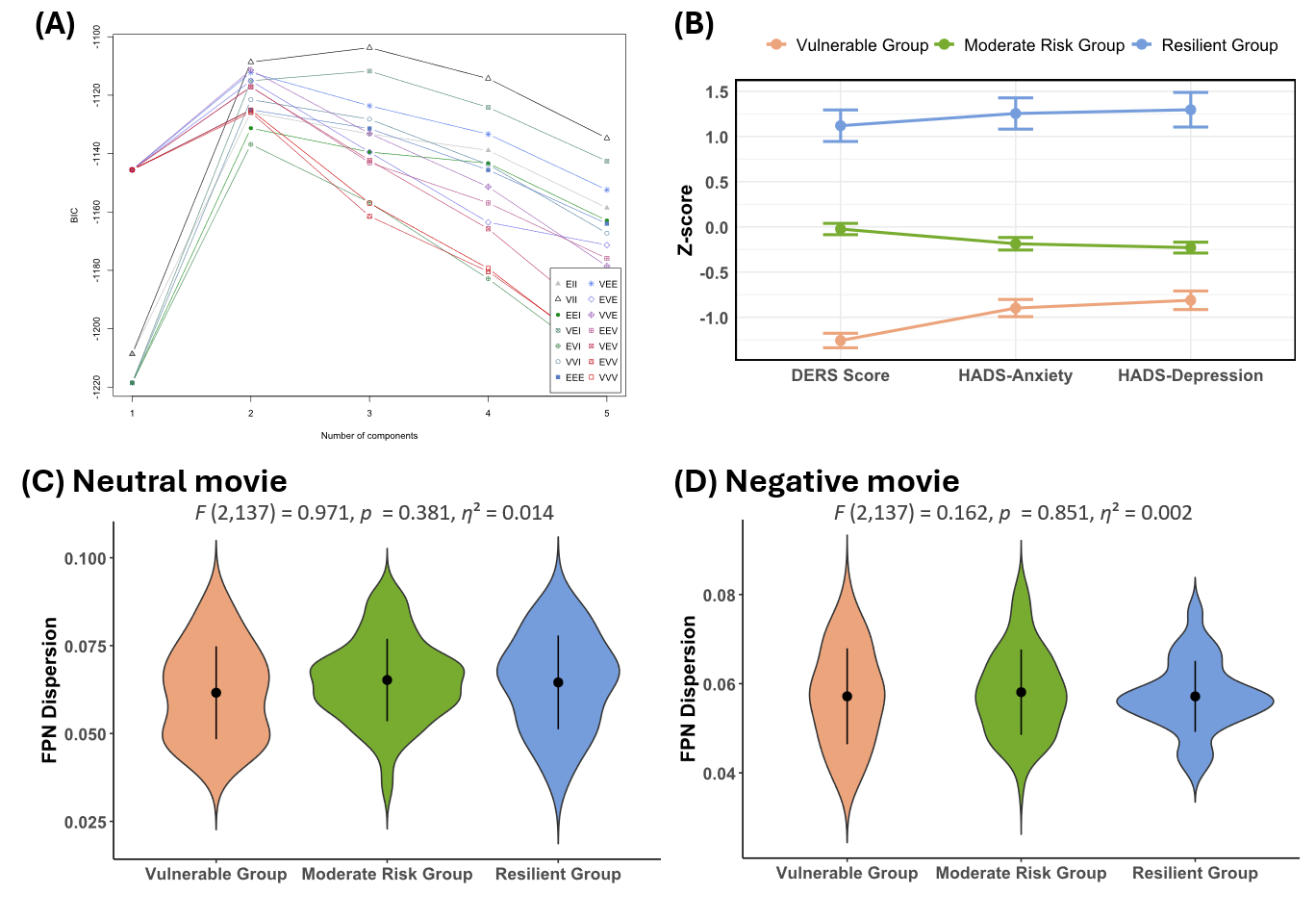

**Figure S9. Three subgroups were identified based on age-adjusted mental health profiles across the whole sample (N=140), and their differences in FPN dispersion. (A)** The best-fitting model was identified as a VII (spherical, varying volume) model with 3 mixture components (BIC = -1103.72). **(B)** Latent profile analysis identified three distinct subgroups with different mental health profiles (i.e., Vulnerable group [N=31], Moderate Risk group [N=83], and Resilient group[N=26]). **(C&D)** No significant differences in FPN dispersion among three groups were observed for both neutral and negative movie conditions.

**Supplementary Tables**

**Table S1. Between-Group Comparisons between resilient and vulnerable subgroups: normality tests, variance homogeneity, and mean differences across conditions**

|  | **Younger adults** | | **Older adults** | |
| --- | --- | --- | --- | --- |
|  | **Neutral**  **FPN dispersion** | **Negative**  **FPN dispersion** | **Neutral**  **FPN dispersion** | **Negative**  **FPN dispersion** |
| **Shapiro-wilk normality test** |  |  |  |  |
| Resilient group | w=0.96 | w=0.97 | w=0.97 | w=0.97 |
| Vulnerable group | w=0.95 | w=0.97 | w=0.94 | w=0.93 |
| **Levene's test** | F(1,70)=2.07 | F(1,70)=1.89 | F(1,66)<0.01 | F(1,66)=0.13 |
| **Welch's t-test** | t=0.04 | t=-0.95 | t=-0.21 | t=3.33** |
| **Mann–Whitney *U* test** | w=569 | w=509 | w=320 | w=531** |
| **Hedges' g** | g=0.01 | g=-0.25 | g=-0.07 | g=1.07 |

Note: **: p<0.01.
